# Deep gastropod relationships resolved

**DOI:** 10.1101/457770

**Authors:** Cunha Tauana Junqueira, Giribet Gonzalo

## Abstract

Gastropod mollusks are arguably the most diverse and abundant animals in the oceans, and are successful colonizers of terrestrial and freshwater environments. Here we resolve deep relationships between the five major gastropod lineages - Caenogastropoda, Heterobranchia, Neritimorpha, Patellogastropoda and Vetigastropoda - with highly congruent and supported phylogenomic analyses. We expand taxon sampling for underrepresented lineages with new transcriptomes, and conduct analyses accounting for the most pervasive sources of systematic errors in large datasets, namely compositional heterogeneity, site heterogeneity, heterotachy, variation in evolutionary rates among genes, matrix completeness and gene tree conflict. We find that vetigastropods and patellogastropods are sister taxa, and that neritimorphs are the sister group to caenogastropods and heterobranchs. With this topology, we reject the traditional Archaeogastropoda, which united neritimorphs, vetigastropods and patellogastropods, and is still used in the organization of collections of many natural history museums. Several traits related to development and life history support our molecular results. Importantly, the time of differentiation of the embryonic 4d cell (mesentoblast, responsible for mesoderm formation), differs between the two major clades, highlighting the degree of conservation and significance of development in the evolution of gastropods, as it is also known for spiralian animals more broadly.

## 1 Introduction

Gastropods are one of the most diverse clades of marine animals [1], and the only mollusk group to successfully colonize terrestrial environments. Besides the number of species, which is in the order of many tens of thousands considering just the described extant diversity, gastropods have a high degree of morphological disparity - snails, limpets and slugs with complex patterns in shell shape, coloration, sizes - and inhabit all kinds of environments and depths. Gastropods have spiral embryo cleavage, an array of developmental modes (direct and indirect, with more than one type of larva), and undergo torsion of the body during development. Five main lineages are currently recognized: Caenogastropoda (e.g., cowries, whelks, conchs, cones), Heterobranchia (e.g., bubble snails, nudibranchs, most terrestrial snails and slugs), Neritimorpha (nerites), Patellogastropoda (true limpets), and Vetigastropoda (e.g., abalones, keyhole limpets, turban snails, top shells).

Early classifications included members of the vetigastropods, patellogastropods and neritimorphs in the Archaeogastropoda [2, 3]. With the first numerical cladistic analysis of morphological data, patellogastropods were recovered as the sister group to all other gastropods, which were united in the clade Orthogastropoda [4, 5]. The sister group relationship of the most diverse lineages, the heterobranchs and caenogastropods into the clade Apogastropoda, has been consistently recovered in most morphological and molecular analyses, but other than that, almost all possible topologies for gastropod relationships have been proposed [for a historical review, see 6], with early molecular studies having mixed success in recovering even the well-established monophyly of gastropods or some of the main lineages [7–11]. The first transcriptomic analyses of the group were able to reject several hypotheses, including the clade Orthogastropoda [12]. However, different methods still resulted in contrasting topologies, and three hypotheses remain [12]. The major uncertainty is the position of Neritimorpha, which is recovered either as the sister group to Apogastropoda, or as the sister group to a clade of Patellogastropoda and Vetigastropoda, in this case forming the traditional Archaeogastropoda. The third remaining hypothesis has vetigastropods as the sister lineage to all other gastropods [12].

Although the most diverse gastropod lineages were well sampled in the transcriptomic analyses of Zapata et al. [12], the dataset had only one species of Patellogastropoda and two of Neritimorpha, which are crucial for the proper rooting of the gastropod tree. Furthermore, several biases known to be present in large genomic datasets were not accounted for in the phylogenetic methods used so far to resolve gastropod relationships. Heterogeneity in the stationary frequency of amino acids among samples is one such issue that can artificially group taxa that are actually not closely related based on convergent amino acid composition [13]. Within-site rate variation through time (heterotachy) is another likely violation [14]. Some genes with slow rates of evolution (e.g., ribosomal protein genes) have also been shown to bias phylogenetic inference [15, 16], while genes with fast rates and high levels of saturation can cause long-branch attraction [17]. An additional model violation comes from gene tree discordance, not accounted for by concatenation methods, that can be caused by incomplete lineage sorting and be particularly relevant in areas of the tree with short internal branches [18–20], such as the radiation of crown gastropods at the Ordovician [12, 21]. More commonly considered issues include rate heterogeneity between sites and missing data.

Our goal was to resolve between the three hypotheses for the early divergences of gastropods. We present an extended sampling of Neritimorpha and Patellogastropoda by producing new transcriptomes, and complement the dataset with the latest published gastropod transcriptomes and with increased representation for the closest outgroups - bivalves, scaphopods and cephalopods. We build a series of matrices and employ a variety of methods and models to account for the most common and relevant potential sources of systematic error in large datasets, namely compositional heterogeneity, site heterogeneity, heterotachy, variation in evolutionary rates among genes, matrix completeness and gene tree conflict.

## 2 Methods

### 2.1 Sampling and sequencing

We sequenced the transcriptomes of 17 species, mostly patellogastropods and neritimorphs, and combined them with published transcriptome sequences from 39 other gastropods and 18 mollusk outgroups, for a total of 74 terminals. All new data and selected published sequences are paired-end Illumina reads. New samples were fixed in RNA*later* (Invitrogen) or flash frozen in liquid nitrogen. RNA extraction and mRNA isolation were done with the TRIzol Reagent and Dynabeads (Invitrogen). Libraries were prepared with the PrepX RNA-Seq Library kit using the Apollo 324 System (Wafergen). Quality control of mRNA and cDNA was done with a 2100 Bioanalyzer, a 4200 TapeStation (Agilent) and the Kapa Library Quantification kit (Kapa Biosystems). Samples were pooled in equimolar amounts and sequenced in the Illumina HiSeq 2500 platform (paired end, 150 bp) at the Bauer Core Facility at Harvard University. New sequences were deposited in the NCBI Sequence Read Archive (BioProject XXX); voucher information, library indexes and assembly statistics are available in the Supplementary Material.

### 2.2 Transcriptome assembly

Both new and previously published transcriptomes were assembled from scratch with a custom pipeline (full details and scripts in the Supplementary Material). Raw reads were cleaned with RCorrector [22] and Trim Galore! [23], removing unfixable reads (as identified by RCorrector), Wafergen library adapters and reads shorter than 50 bp. Filtered reads were compared against a dataset of mollusk ribosomal RNAs and mitochondrial DNA and removed with Bowtie2 v2.2.9 [24]. This dataset was created from the well-curated databases SILVA [25] (18S and 28S rRNAs), AMIGA [26] (mtDNA) and from GenBank [27] (5S and 5.8S rRNAs). Reads were assemble into transcripts with Trinity v2.3.2 [28, 29] (–SS_lib_type FR for our new strand-specific data generated with Wafergen kits; precise information was not available from published data, so the default non-strand-specific mode was used for reads downloaded from SRA). A second run of Bowtie2 was done on the assemblies, before removing transcripts with sequence identity higher than 95% with CD-HIT-EST v4.6.4 [30, 31]. Transcripts were then translated to amino acids with TransDecoder v3.0 [29], and the longest isoform of each gene was retained with a custom python script (*choose_longest_iso.py*). The completeness of the assemblies was evaluated with BUSCO by comparison with the Metazoa database [32].

### 2.3 Matrix construction

We built four matrices to account for extreme evolutionary rates, amino acid composition heterogeneity and different levels of matrix completeness. Scripts and a detailed pipeline are available in the Supplementary Material. Orthology assignment of the peptide assemblies was done with OMA v2.0 [33]. We then used a custom python script (*selectslice.py*) to select all orthogroups for which at least half of the terminals were represented (50% taxon occupancy), resulting in a matrix with 1059 genes (Matrix 1) (Figure 1). Each orthogroup was aligned with MAFFT v7.309 [34], and the alignment ends were trimmed to remove positions with more than 80% missing data with a custom bash script (*trimEnds.sh*). To avoid possible biases, saturation and long-branch attraction, Matrix 2 was built by removing from Matrix 1 the 20% slowest and the 20% fastest evolving genes, as calculated with TrimAl [35], for a final size of 635 genes (Figure 1). Matrix 3 is the subset of 962 genes from Matrix 1 that are homogeneous regarding amino acid composition. Homogeneity for each gene was determined with a simulation-based test from the python package p4 [13, 36], with a script modified from Laumer et al. [37] (*p4_compo_test.py*) and a conservative p-value of 0.1. Finally, a subset of 149 genes with 70% taxon occupancy constitutes Matrix 4 (Figure 1). For inference methods that require concatenation, genes were concatenated using Phyutility [38]. We further reduced composition heterogeneity in Matrices 1 and 2 by recoding amino acids into the six Dayhoff categories [39] with a custom script (*recdayhoff.sh*).

**Figure1:**
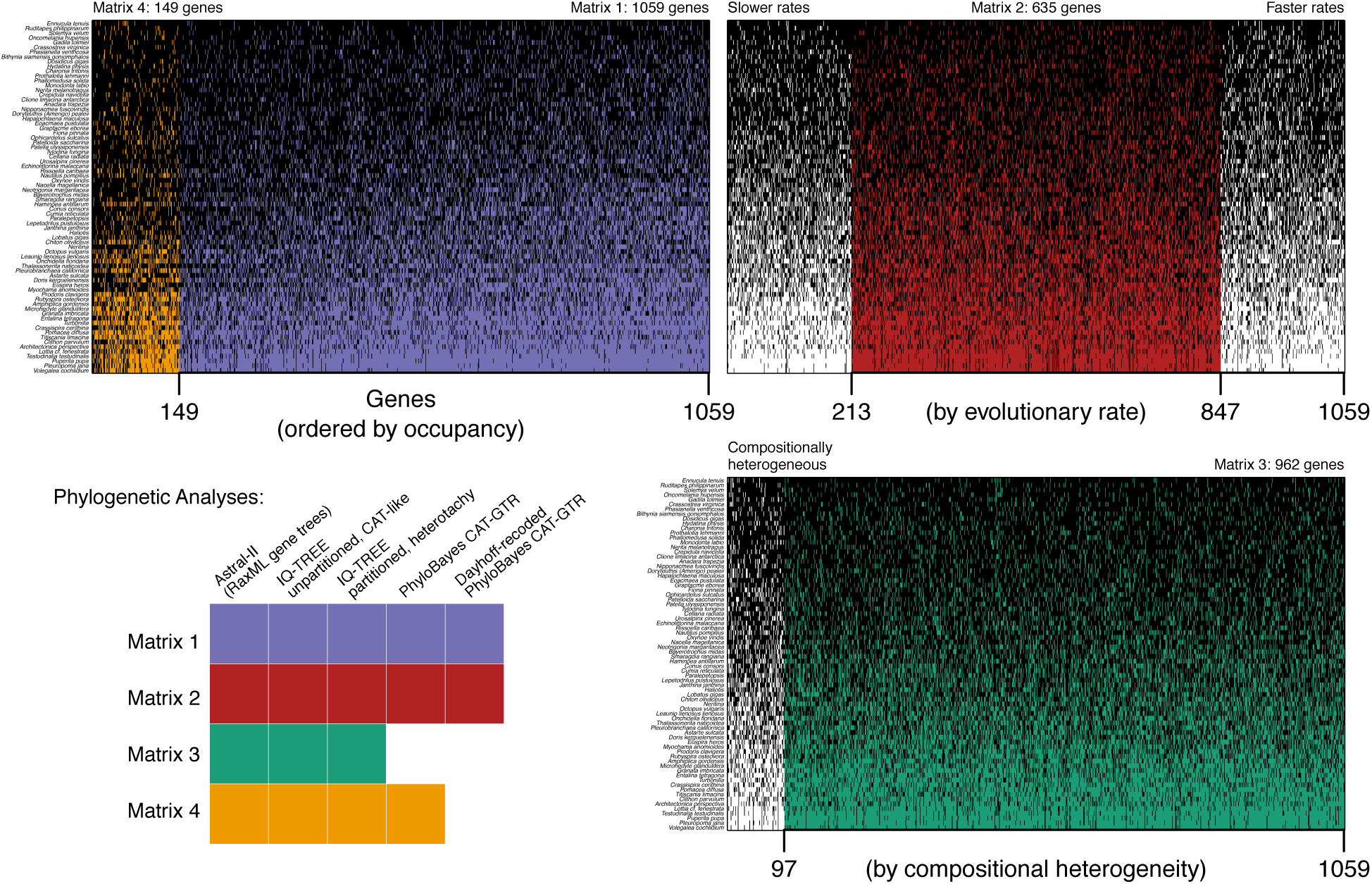
Matrices and phylogenetic methods used to infer gastropod relationships. With 50% taxon occupancy, Matrix 1 is the largest, with 1059 genes. Matrix 4 is the subset of the best sampled 149 genes, with 70% taxon occupancy. Genes and species are sorted with the best sampling on the upper left. Matrix 2 is the subset of 635 genes after ordering all genes by evolutionary rate and removing the 20% slowest and 20% fastest evolving genes. Matrix 3 includes the 962 genes that are homogeneous in amino acid composition; genes are ordered by p-value of the homogeneity test. Black cells indicate genes present for each species. Check Methods for details.

### 2.4 Phylogenetic analyses

Amino acid matrices were used for phylogenetic inference with a coalescent-based approach in Astral-II v4.10.12 [40], with maximum likelihood (ML) in IQ-TREE MPI v1.5.5 [41–43], and with Bayesian inference in PhyloBayes MPI v1.7a [44]. The two Dayhoff-recoded matrices were analyzed in PhyloBayes (Figure 1). Full details and scripts are explained in a custom pipeline in the Supplementary Material. For the coalescent-based method, gene trees were inferred with RAxML v8.2.10 [45] (-N 10-m PROTGAMMALG4X) and then used as input for Astral-II for species tree estimation. For each concatenated matrix, we inferred the best ML tree with two strategies: a gene-partitioned analysis with model search including LG4 mixture models and accounting for heterotachy (-bb 1500-sp partition_file-m MFP+MERGE-rcluster 10-madd LG4M, LG4X-mrate G,R,E); and a non-partitioned analysis with model search also including the C10 to C60-profile mixture models [46] (ML variants of the Bayesian CAT model [47]) (-bb 1500-m MFP+MERGE-rcluster 10-madd LG4M,LG4X,LG+C10,LG+C20,LG+C30,LG+C40,LG+C50,LG+C60-mrate G,R,E). The search for the models LG+C60 (Matrices 1 and 3) and LG+C50 (Matrix 1) required more memory than available, and these models were disregarded for the respective matrices. PhyloBayes was run with the CAT-GTR model on a subset of the concatenated alignments (matrices 1, 2 and 4), discarding constant sites to speed up computation.

## 3 Results and discussion

### 3.1 Main gastropod relationships

Our main goal was to resolve the deep nodes of the gastropod tree and distinguish between three hypotheses of the relationships among its five main lineages. All but one of our inference methods and matrices congruently support a clade uniting Vetigastropoda and Patellogastropoda, and Neritimorpha as the sister group to Apogastropoda (Figure 2). The only exception is the coalescent-based analysis on the smallest dataset of 149 genes (Astral, Matrix 4), in which these two key nodes were left unresolved (all tree files are available in the Supplementary Material). Accordingly, the few analyses with lower support on these nodes also refer to the smaller Matrix 4, which is unsurprising given that it comprises fewer informative sites in concatenated analyses, and less genes in the coalescent-based analysis [48]. In summary, the resulting topology is congruent based on an array of analyses testing for the major common sources of systematic error in phylogenomic datasets, including gene tree discordance, compositional heterogeneity, heterotachy, site heterogeneity, variation in evolutionary rates, and missing data.

**Figure 2:**
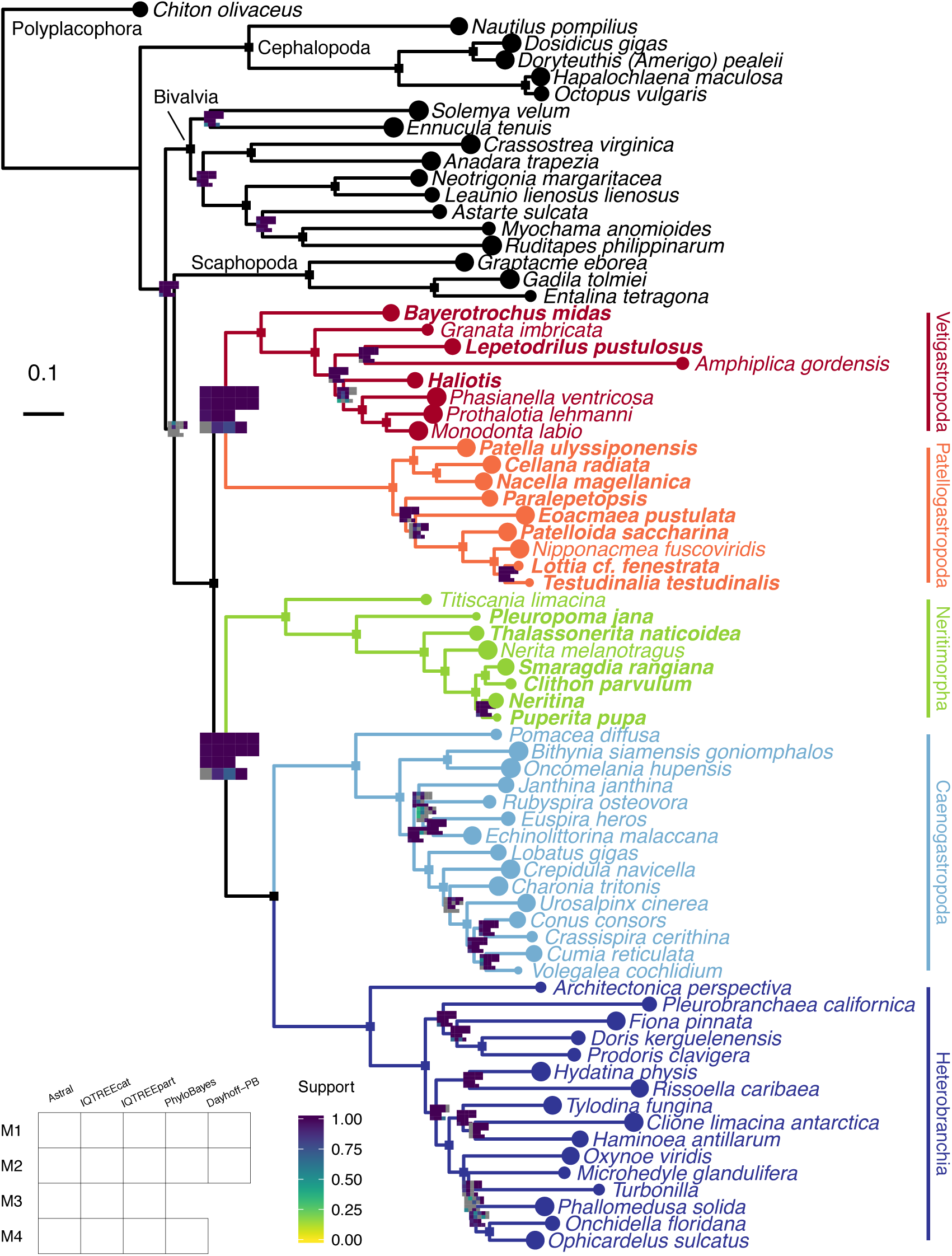
Gastropod phylogeny inferred from the largest matrix (M1) with Maximum Likelihood and a profile mixture model (IQTREEcat). A single square marks branches where all analyses had full support; branches where at least one analysis had less than full support are marked with a plot, colored in a continuous scale according to support value, from 0 to 1. Grey squares in the plots represent splits that were absent in a given analysis. M1-M4: Matrices 1-4; IQTREEpart: ML partitioned analysis; Dayhoff-PB: Bayesian analysis on a matrix recoded according to the 6 Dayhoff categories. Check Methods for details.

This exact topology for gastropod relationships has been previously recovered by a few molecular [12, 49] and total evidence [6] analyses, with numerous alternatives proposed in the literature [e.g. 5, 6, 10, 49–xs51], even within the same studies. With 17 analyses (combinations of four subsampled matrices, two data types - amino acids and Dayhoff recoding, and four inference methods/models), for the first time we find strong congruence and support for the tree presented in Figure 2. With that we reject the clade Archaeogastropoda, proposed almost a century ago by Thiele [2], which united Neritimorpha, Vetigastropoda and Patellogastropoda. Although this grouping had given way to other predominant hypotheses along the years (e.g., Eogastropoga *vs.* Orthogastropoda divergence), this classification is still used in the organization of malacology and paleontological collections of many natural history museums.

We find strong support for the position of neritimorphs as the sister group to apogastropods, a relationship that is further supported by important developmental characters [52, 53]. Vetigastropods and patellogastropods differ from neritimorphs and apogastropods in the time of differentiation of the 4d cell (mesentoblast), a key embryonic cell that gives rise to the mesoderm in spiralians [54–56]. In Vetigastropoda and Patellogastropoda, the mesentoblast is formed at the 63-cell stage; in the other gastropod lineages, formation of the mesentoblast is accelerated, happening sometime between the 24- and 48-cell stages depending on the species [52, 53]. Given the highly conserved nature of the early spiral cleavage program and of cell fates across spiralian phyla [54–56], the congruence between our molecular results and the variation in development highlights the significance of such traits also for the evolution of the main gastropod lineages. A few other reproduction and life history traits distinguish the two main clades of gastropods recovered here. While vetigastropods and patellogastropods are mostly broadcast spawners, neritimorphs and apogastropods have internal fertilization and often complex reproductive behaviors and anatomy [57]. For these latter groups, eggs are usually encapsulated, with either direct development or the release of a feeding veliger larva in the water, while embryos of vetigastropods and patellogastropods first develop into a non-feeding trochophore larva in the plankton [58]. In addition, neritimorphs and apogastropods have invaded freshwater and terrestrial environments several times. All of these traits are likely connected, but order of appearance and causal relations remain to be investigated.

Important questions that remain regarding major gastropod relationships include the position of Cocculiniformia, Neomphalina and the hot-vent taxa, smaller deep sea clades that have been considered somehow related to vetigatropods, neritimorphs, patellogastropods, or as independent branches in the gastropod tree. They are yet to be sampled in a phylogenomic analyses.

Regarding overall mollusk relationships, we recover a well supported clade of gastropods, bivalves and scaphopods in all analyses; however, as in previous phylogenomic efforts [59, 60], relationships between these three groups are unstable (Figure 2). The Dayhoff datasets and most of the ML analyses with the profile mixture model result in a clade of gastropods and scaphopods; while most coalescent-based trees recover a clade of bivalves and scaphopods; and finally, the ML partitioned analyses produce a clade of gastropods and bivalves. Perhaps a way ahead to resolve such hard nodes will be to use other types of data, such as genomic rearrangements and presence/absence of genes from complete genomes.

### 3.2 A note about convergence in PhyloBayes

While PhyloBayes converged on Dayhoff-recoded datasets, analyses on the more complex amino acid matrices did not converge for all parameters. The problem was especially pronounced for the large matrices (a summary table for all analyses is given in the Supplementary Material). We observed that convergence issues were mostly due to small differences between chains regarding the position of one or few derived terminals within the outgroups or within apogastropods, whose relationships were not the goal of this study. We suspect this may be caused by a problem in topology proposals for these derived nodes, leading some of the chains to get stuck in local maxima. One example comes from the Dayhoff analysis of Matrix 1: the initial two chains seemed to be very far from topological convergence (maxdiff=1) even after more than 20,000 generations. Upon closer inspection, both trees were basically indistinguishable, with the only variation being the position of *Charonia* or *Crepidula* as the sister group to Neogastropoda. Removal of either one of the two terminals from the treelist file with a custom script (*remove_terminal.py*) resulted in the same converged topology (tree files in the Supplementary Material). For that particular analysis, we ran two additional independent chains that converged without presenting this issue. This behavior was recently discussed [61], and perhaps has been underreported in the literature.

### 3.3 Relationships within gastropod lineages

This is the first genomic-scale dataset for Patellogastropoda and Neritimorpha. Internal relationships of patellogastropods have presented incongruent results even among studies using the same type of data (reviewed in Lindberg [62] and Nakano & Sasaki [63]). We consistently recover Nacellidae (*Cellana*, *Nacella*) as the sister group of Patellidae (Figure 2), a clade originally supported by some of the earliest morphological [64 and mitochondrial phylogenies [65]. Nacellids have also been placed either as a grade at the base of the tree [66] or closer to Lottiidae [67], and the current taxonomic classification has Nacellidae in the superfamily Lottioidea [68]; our results indicate the family should be transferred to Patelloidea. Another interesting finding regards *Eoacmaea*, which had gained family and superfamily status due to being recovered as the sister taxon to all other patellogastropods with mitochondrial markers [67]. None of our results recover this position, but rather indicate that the genus is either part of Lottiidae (most ML and Bayesian results), which was its original assignment, or is sister group to the Lottioidea families Neolepetopsidae (*Paralepetopsis*) and Lottiidae (*Patelloida*, *Nipponacmea*, *Lottia*, *Testudinalia*) (coalescent-based trees and one ML tree) (Figure 2).

Neritimorphs had mostly congruent phylogenies recovered from 28S rRNA data [69] and mitogenomes [70]. Our reconstruction supports the same topology, with Neritopsoidea (*Titiscania*) as the sister group to all other neritimorphs, followed by the divergence between Helicinoidea (*Pleuropoma*) and Neritoidea. Within the latter, we also recover a monophyletic Neritidae as the sister group of Phenacolepadidae (*Thalassonerita*). The nested position of *Smaragdia* inside Neritininae disagrees with its current classification in its own subfamily [68].

Vetigastropoda and Heterobranchia had similar taxon representation as in Zapata et al. [12] (restricted to only transcriptomes with high sequencing quality, and with newly sequenced replacements for some vetigastropod families). As expected, the relationships are the same, and highlight the need for future studies focused on each group, given the uncertain position of *Haliotis* in Vetigastropoda, and low resolution of internal relationships of panpulmonates in Heterobranchia.

We substantially increased sampling of Caenogastropoda by adding the latest published transcriptomes of eight families. Despite that, caenogastropods are the most diverse gastropod lineage, with over a hundred families, and the following results are still limited in sampling. We recover a monophyletic Neogastropoda; its internal relationships differ from a molecular study with denser taxon sampling [71], in that we find Buccinoidea (*Cumia*, *Volegalea*) closer to Conoidea (*Conus*, *Crassispira*) than to Muricoidea (*Urosalpinx*). We also recover a monophyletic Truncatelloidea (*Bithynia*, *Oncomelania*) as the sister group to all other Hypsogastropoda. The relative position of Tonnoidea *(Charonia)* and Calyptraeoidea (*Crepidula*) regarding Neogastropoda is unclear, nonetheless, this close relationship between them, and also of Stromboidea (*Lobatus*), agrees with previous molecular studies [71, 72]. The branching pattern of the closest relatives of neogastropods reveal a paraphyletic Littorinimorpha [68].

## 5 Authors’ contributions

TJC and GG conceived the study, collected and identified specimens. TJC carried out lab work, analyzed the data and drafted the manuscript. Both authors improved the manuscript and gave final approval for publication.

## Acknowledgments

We thank James Reimer for kindly providing support for fieldwork. We also thank Vanessa Knutson, Shawn Miller, Hin Boo Wee, Kristen Soong and Taku Ohara for help in the field. Don Colgan (Australian Museum) and Lluis Cardona (University of Barcelona) each donated a specimen. The computations in this paper were done on the Odyssey cluster supported by the FAS Division of Science, Research Computing Group at Harvard University. We thank software developers and community forums for help with software, in particular Brian Haas (Trinity), Adrian Altenhoff (OMA), and Bui Quang Minh (IQTREE). We thank Bruno de Medeiros for discussions and help with scripts.

## Supplementary material and data access

New transcriptomes were deposited in the NCBI Sequence Read Archive (BioProject XXX). Supplementary materials include detailed pipelines, tree files, images and tables, deposited in Harvard Dataverse doi:XXX. Scripts are available on github.com/tauanajc/phylo_scripts.

